# More than efficacy revealed by single-cell analysis of antiviral therapeutics

**DOI:** 10.1101/606715

**Authors:** Wu Liu, Mehmet U. Caglar, Zhangming Mao, Andrew Woodman, Jamie J. Arnold, Claus O. Wilke, Craig E. Cameron

## Abstract

Development of antiviral therapeutics emphasizes minimization of the effective dose and maximization of the toxic dose, first in cell culture and later in animal models. Long-term success of an antiviral therapeutic is determined not only by its efficacy but also by the duration of time required for drug-resistance to evolve. We have developed a microfluidic device comprised of ~6000 wells, with each well containing a microstructure to capture single cells. We have used this device to characterize enterovirus inhibitors with distinct mechanisms of action. In contrast to population methods, single-cell analysis reveals that each class of inhibitor interferes with the viral infection cycle in a manner that can be distinguished by principal component analysis. Single-cell analysis of antiviral candidates reveals not only efficacy but also properties of the members of the viral population most sensitive to the drug, the stage of the lifecycle most affected by the drug, and perhaps even if the drug targets an interaction of the virus with its host.

## INTRODUCTION

Over the past few decades, the world has witnessed outbreaks of myriad RNA viruses, including West Nile Virus, severe acute respiratory syndrome (SARS) coronavirus, chikungunya virus, Ebola virus, Zika virus, and most recently the poliovirus (PV)-related viruses: enterovirus D68 (EV-D68) and enterovirus A71 (EV-A71) (Antona et al., 2016; Baize et al., 2014; Burt et al., 2012; Campbell et al., 2002; Campos et al., 2015; Peiris et al., 2003). Unfortunately, viral emergence has outpaced the discovery and development of compounds capable of treating these pathogens. Because sporadic outbreaks come, go, and may never come again, development of broad-spectrum therapeutics exhibiting high barriers to resistance would have the greatest value. Unbiased screening of chemical libraries for antiviral agents using cell-based assays have no problem identifying active compounds of high potency. However, identifying the target and predicting the likelihood for evolution of resistance generally takes years of effort following compound discovery.

As a part of a study evaluating PV infection dynamics on the single-cell level, we observed that a chain-terminating antiviral ribonucleotide selectively eliminates the most-fit members of the viral population (Guo et al., 2017). This class of antiviral agent has always been touted as having a high barrier to resistance (Jordheim et al., 2013). The typical explanation for this high barrier is that amino acid substitutions in the active site of the viral RNA polymerase conferring resistance to the antiviral ribonucleoside also impair the specificity and/or efficiency of incorporation of natural ribonucleotides (Carroll et al., 2003). Elimination of the most-fit members of the viral population by an antiviral agent requires that resistance emerge from the surviving, low-fitness member of the population, which would ultimately require restoration of fitness for the population to survive the myriad mechanisms of host restriction (Andino and Domingo, 2015). Restoration of fitness may be the insurmountable barrier precluding the development of resistance to chain-terminating antiviral ribonucleotides.

This study was designed to determine the extent to which single-cell analysis of antiviral agents can contribute to our understanding of antiviral therapeutics relative to traditional approaches. We describe a microfluidics device that can be used to produce complete dose-response curves. We have used this device to compare three, mechanistically-distinct classes of antiviral agents: a PV polymerase inhibitor (2’-*C*-methyl-adenosine, 2’-*C*-Me-A); a PV protease inhibitor (rupintrivir); and two HSP90 inhibitors (geldanamycin, GA, and ganetespib, GS). We find that single-cell analysis distinguishes these classes of inhibitors. We suggest that addition of single-cell analysis to the existing paradigm for preclinical development of antiviral therapies may have the potential to identify leads with limited potential for development of resistance.

## RESULTS

### A device for single-cell analysis of the activity of antiviral agents

Our initial foray into single-cell analysis of PV infection dynamics used cell density to control single-cell occupancy of wells of a microfluidic device (Guo et al., 2017). What this means practically is that most wells are empty. Reducing the number of events monitored by a mere factor of two takes us below the number of events required for statistical analysis of the data. Acquisition of a dose-response curve was therefore impossible using this first-generation device.

To address this problem, we created a multi-layer device as previously described (**Fig. 1a**) (Guo et al., 2017) but added a trapping structure to the channel layer of the device (**Figs. 1b** and **S1a**). Position of the trapping structure was guided by simulation (**Fig. 1c**), leading to placement of the trapping structure on the rear wall of each well (**Fig. 1d**). The device contains five independent zones, each containing 1140 wells for a total of 5700 wells. The device mounts easily to the stage of a microscope (**Fig. S1b**); the device in cross-section is shown in **Fig. S2**. Single-cell occupancy of the device was sensitive to the width of the channels used for loading (**Fig. S3a**) and cell density (**Fig. S3b**) with maximum occupancy near 90% (**Figs. 1d** and **S3**). Under conditions in which an inhibitor reduces infection to 10%, ~100 infected cells will be observed using this device. This number of events is more than enough for statistical analysis.

**Figure 1.**
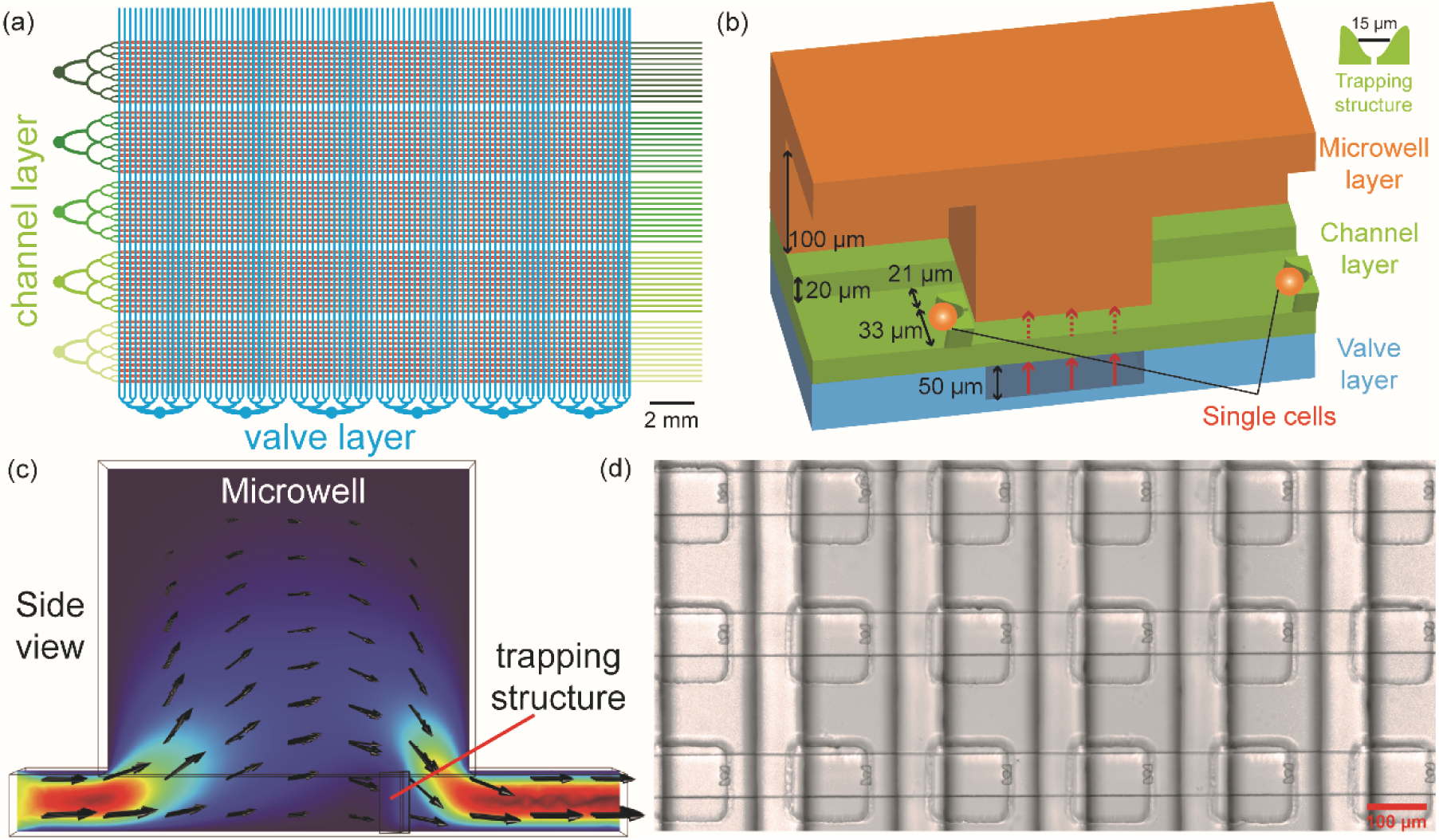
Addition of cell-trapping microstructures to wells of a microfluidic device enhances single-cell occupancy. (**a**) Schematic of the device. The following layers exist: microwell (not shown explicitly); channel (green); and valve (cyan). The device is divided into five sections of 1140 microwells (different shades of green for the channel layer), for a total of 5700 microwells. (**b**) Schematic of two wells of the device with all relevant dimensions indicated. The microwell layer creates a physical barrier between adjacent wells. The barrier between adjacent wells is sealed with water emanating from the valve layer. Water in valve layer is sealed by application of air under pressure (20-30 psi). (**c**) Simulation of the flow velocity field in a microwell indicated that the outlet would maximize cell trapping by the microstructure. (**d**) Image showing cells captured by the microstructures in the microwells. The device was infused with cells (5 × 10^5^ cells/mL) at rate of 0.5 μL/min. Under optimal conditions, 86.1% (4902) microwells of the device contained single cells.

### Re-evaluation of a PV polymerase inhibitor

Viral polymerases represent a well-established target of antiviral therapeutics. For example, cocktails used to treat human immunodeficiency virus infection and hepatitis C virus (HCV) infection include synthetic nucleoside analogues targeting reverse transcriptase and RNA-dependent RNA polymerase (RdRp), respectively (Cihlar and Ray, 2010; Sofia et al., 2012). The non-obligate, chain-terminating antiviral ribonucleoside, 2’-*C*-Me-A, is the prototype for the HCV RdRp inhibitor, sofosbuvir and related compounds (Eltahla et al., 2015). In a previous study, we evaluated infections that survived treatment when present at a concentration that reduces the number of infections by 50% (IC50). To our surprise, we observed selective ablation of cells infected with virus variants capable of the fastest rates of replication and the highest yields of replicated RNA (Guo et al., 2017). This experiment suffered from the inability to evaluate concentrations of drug higher than the value of the IC50. Here, we use 2’-*C*-Me-A to validate the new device.

For all experiments reported herein, we infect HeLa S3 cells off chip with PV engineered to express enhanced green fluorescent protein (PV-eGFP) at a multiplicity of infection of 0.5 plaque-forming units per cell (PFU/cell), pellet and wash cells to remove free virus, suspend cells to a density of 5 × 10^5^ cells/mL, and load the microfluidic device. Each well is monitored for eGFP fluorescence every 30 min for a 24 h period, which gives rise to a population of time courses exhibiting substantial between-cell variability (**Fig. S4a**). We plot the percentage of infected cells as a function of drug concentration and fit to a hyperbola to obtain a value for the IC50 value. In the case of 2’-*C*-Me-A, concentrations greater than 50 µM exhibit toxicity (Stuyver et al., 2006); therefore, these data do not fit to a hyperbola (**Fig. S4b**). It is clear from this experiment, however, that the approach is sensitive enough to acquire data for a complete dose-response analysis (**Fig S4b**).

A major advantage of the single-cell platform is that the effect of a drug on the viable population can be determined. To analyze the single-cell data, we use five phenomenological parameters extracted from each time course (**Fig. S4c**). These parameters are: *maximum* fluorescence observed, which correlates to yield of genomic RNA; *slope* at the time of half-maximum fluorescence, which correlates to replication speed; *infection time*, which, to a first approximation, is the time it takes for an infection to go from start to finish akin to the virus generation time; *start-point*, which is the earliest time in which fluorescence can be detected; and *midpoint*, the time of half-maximum fluorescence. This analysis leads to a distribution of values formed from values measured in each cell. Our statistical analysis uses an unpaired two-tailed t-test to determine if a significant difference exists for the means of a given parameter under two experimental conditions. In these experiments, the area under the curve defining the distribution reflects the number of events monitored. We do not attempt to interpret a difference in the “fine” structure of the distributions.

Using this data-analysis pipeline to evaluate outcomes in the absence and presence of 2’- *C*-Me-A (50 µM), we observed a statistically significant difference in the mean values for the maximum (**Fig. S4d**) and slope (**Fig. S4e**) parameters without a statistically significant change in the mean values for the remaining parameters (**Figs. S4f** - **S4h**). Tabulation of the data and statistical analysis for this experiment is provided in **Tables S1** and **S2**. These observations reproduce outcomes presented by us previously using the first-generation device (Guo et al., 2017). Members of the viral population with highest values for maximum and slope parameters are most sensitive to 2’-*C*-Me-A.

### Evaluation of a PV 3C-protease inhibitor

The observation that a subset of the viral population was most sensitive to 2’-*C*-Me-A was unexpected but could be easily rationalized by the notion that increased replication efficiency should correlate to increased 2’-*C*-Me-AMP incorporation efficiency. Do inhibitors of other viral enzymes exhibit the same selectivity? Proteases of retroviruses and positive-sense RNA viruses responsible for polyprotein processing represent a second category of well-established antiviral therapeutics used in the cocktails to treat HIV and HCV, respectively (de Leuw and Stephan, 2018; Pokorna et al., 2009; Wlodawer and Vondrasek, 1998). Inhibitors of the picornaviral protease responsible for polyprotein processing (3C) have been pursued for many years (Wang and Liang, 2010). For this study, we selected rupintrivir, which targets the 3C protease activity of all three types of polioviruses as well as that of other enteroviruses (Binford et al., 2005; De Palma et al., 2008).

We measured the impact of rupintrivir on establishment of PV infection in HeLa S3 cells (**Fig. 2a**). The IC50 value was 8.0 ± 4.0 nM, which agrees with values (20 nM) measured using conventional plaque assays (De Palma et al., 2008). We analyzed the single-cell data obtained at all concentrations (**Tables S3** and **S4**) but focused only on the concentrations approximating the IC50 and 2 × IC50, 10 nM and 20 nM, respectively, to emphasize trends in the data. At both concentrations, we observed a statistically significant difference compared to the control for all parameters: maximum (**Fig. 2b**); slope (**Fig. 2c**); infection time (**Fig. 2d**); start point (**Fig. 2e**); and midpoint (**Fig. 2f**). This outcome with rupintrivir is clearly distinct from that above with 2’- *C*-Me-A. We conclude that not all antiviral agents interfere with the viral population in the same manner or at the same stage(s) of the lifecycle.

**Figure 2.**
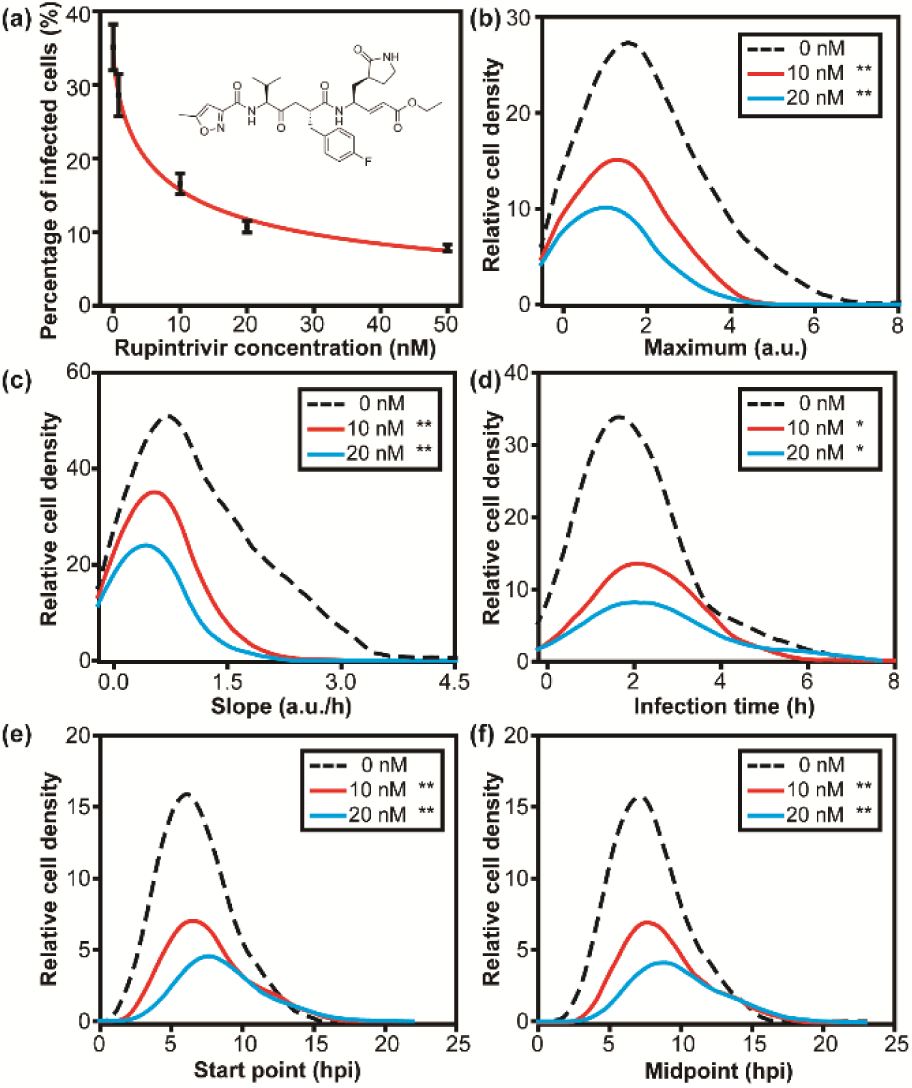
Evaluation of rupintrivir, a PV 3C protease inhibitor. (**a**) Dose-response analysis. Percentage of single, infected (green) cells was determined as a function of rupintrivir concentration. (**b**-**f**) Distributions for each parameter in the presence of 10 or 20 nM rupintrivir were compared to that in the absence of drug using a t-test. A single asterisk indicates a p-value less than 0.05; two asterisks indicate a p-value is less than 0.005. Numerical values for experimental parameters are provided in **Table S3** and statistical analysis in **Table S4**. The parameters presented in the panels are as follows: (**b**) maximum; (**c**) slope; (**d**) infection time; (**e**) start point; (**f**) midpoint.

### Evaluation of HSP90 inhibitors

Compounds antagonizing the function of cellular chaperones represent an emergent class of anti-cancer and antiviral therapeutics (Chatterjee and Burns, 2017; Geller et al., 2012). Chaperones of the heat-shock-protein-90 (HSP90) family are required when the cell is growing fast and/or producing high levels of proteins, as observed in cancer cells or in virus-infected cells. Seminal studies from the Frydman and Andino laboratories demonstrate a clear, essential role for HSP90 in production of infectious PV virions. However, every step of the virus lifecycle has been suggested as a target for HSP90 when viruses outside of the picornavirus family are considered (Geller et al., 2012; Wang et al., 2017). The benefit of targeting antiviral therapeutics to cellular proteins is that the likelihood for evolution of resistance is minimized (Geller et al., 2007).

We have used geldanamycin (GA), because it is a potent inhibitor of HSP90 (nM range) that is active against PV (Geller et al., 2007). The presence of GA reduced the number of infections established by 22 ± 4% with an IC50 value of 30 ± 8 nM (**Fig. 3a**). An earlier study did not observe an impact of GA prior to virus assembly; however, it is possible that a reduction of the magnitude shown here would be concealed by experimental error (Geller et al., 2007). Because infection is monitored by eGFP, which requires virus entry, genome replication, and genome translation, interference with any of these steps would lead to a reduction in the number of eGFP-positive cells. It is clear, however, that translation and folding of eGFP are not altered in the presence of the highest concentration of GA used in this experiment (**Fig. S5**).

**Figure 3.**
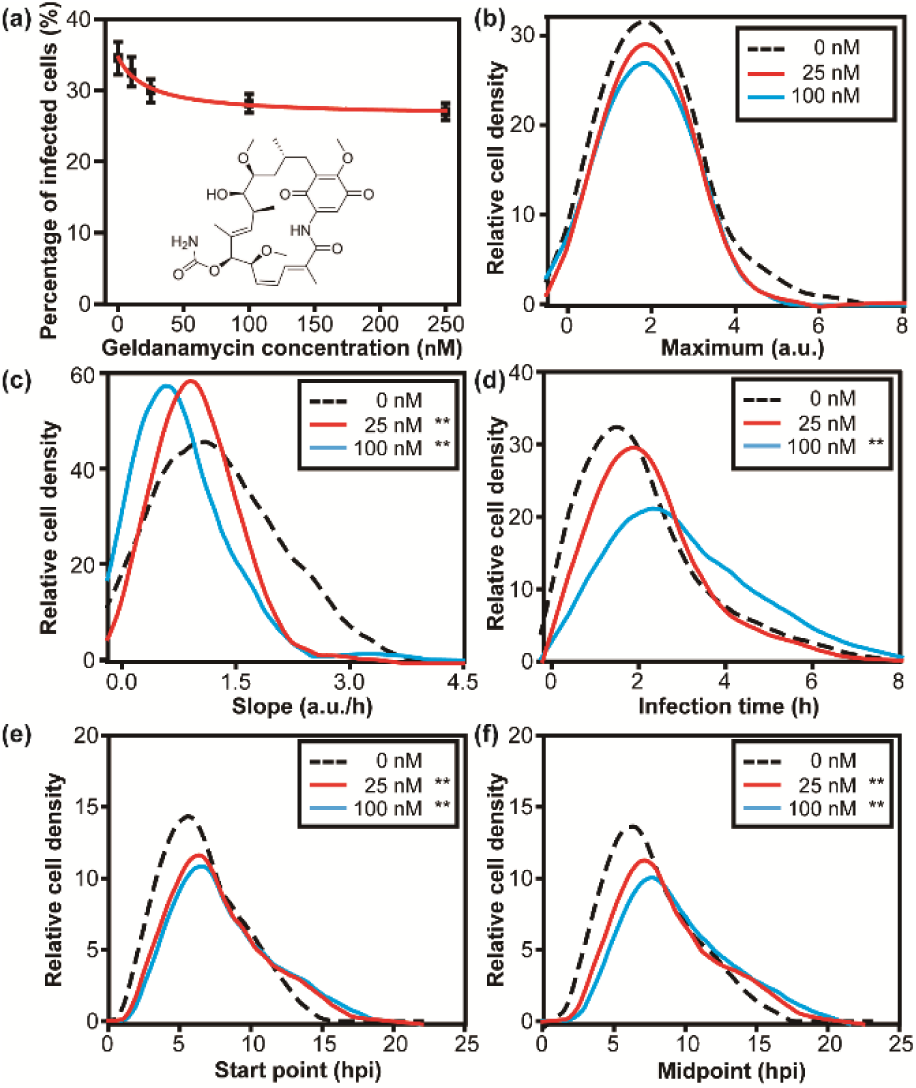
Evaluation of geldanamycin (GA), an HSP90 inhibitor. (**a**) Dose-response analysis. Percentage of single, infected (green) cells was determined as a function of GA concentration. (**b**- **f**) Distributions for each parameter in the presence of 25 or 100 nM GA were compared to that in the absence of drug using a t-test. A single asterisk indicates a p-value less than 0.05; two asterisks indicate a p-value less than 0.005. Numerical values for experimental parameters are provided in **Table S5** and statistical analysis in **Table S6**. The parameters presented in the panels are as follows: (**b**) maximum; (**c**) slope; (**d**) infection time; (**e**) start point; (**f**) midpoint.

Analysis of the single-cell data is presented in **Tables S5** and **S6**. The mean of the distribution of values for the maximum parameter did not change in the presence of GA (**Fig. 3b**), in contrast to the inhibitors targeting viral proteins. Observation of a statistically significant difference in the distribution of the values for the infection-time parameter was concentration dependent (**Fig. 3d**). A statistically significant difference for mean of the distributions for the remaining parameters was observed at concentrations corresponding to the IC50 and above (**Figs. 3c, 3e** and **3f**). A third signature of antiviral action is therefore revealed with GA.

Given the interest in using HSP90 inhibitors as therapeutics for cancer, a variety of compounds exist (Hwang et al., 2009; Sidera and Patsavoudi, 2014). For example, ganetespib (GS) is an HSP90 inhibitor that differs from GA substantially in chemical structure (compare inset in **Fig. S6** to that in **Fig. 3a**), has a higher affinity for HSP90 than GA (Prince et al., 2015), but is thought to have the same mechanism of action (Roe et al., 1999; Ying et al., 2012). Use of GS provides an important test of the capacity of the single-cell analysis to reveal common signature for compounds with common mechanisms.

In contrast to observations with GA, GS reduced the number of infections established by 70 ± 14% with an IC50 of 3 ± 1 nM. Analysis of the single-cell data is presented in **Tables S7** and **S8**. The impact of GS on the five phenomenological parameters were the same as observed for GA (**Fig. S6**). In this case, it is clear that mechanistically identical compounds yield identical signatures at the single-cell level.

### Evaluation of antiviral drug combinations

The long-term utility of an antiviral therapy is determined, in part, by the time required for resistance mutants to emerge. One approach to delaying or even eliminating the emergence of drug-resistant mutants is the use of antiviral combinations (Dunning et al., 2014; Hofmann et al., 2009). What is often evaluated is the ability of two drugs to exhibit greater efficacy (synergy) in preventing infection in combination than observed when either drug is used alone. We evaluated the ability of 2’-*C*-Me-A to synergize with GA by treating cells with the concentrations of one or both at their IC50 values (**Fig. 4a**). In the absence of drug, 34 ± 3% of cells in the device were infected. That number was reduced to 18 ± 2%, 30 ± 3%, or 15 ± 3% in the presence of 2’-*C*-Me-A, GA, or the combination thereof, respectively. Based on this observation alone, the conclusion would be that this combination of antiviral agents is not even additive. Analysis of the entire single-cell dataset is presented in **Tables S9** and **S10**. Because 2’-*C*-Me-A exhibits the most substantial antiviral effect relative to the DMS0 control, here we compare the combination to 2’-*C*-Me-A alone. We observed a statistically significant difference for all parameters (**Figs. 4b - 4f**). Single-cell analysis therefore has the ability to reveal efficacy of drug combinations masked at the population level. We performed the comparable experiment with GS and reached the same conclusion (**Fig. S7, Tables S10** and **S11**).

**Figure 4.**
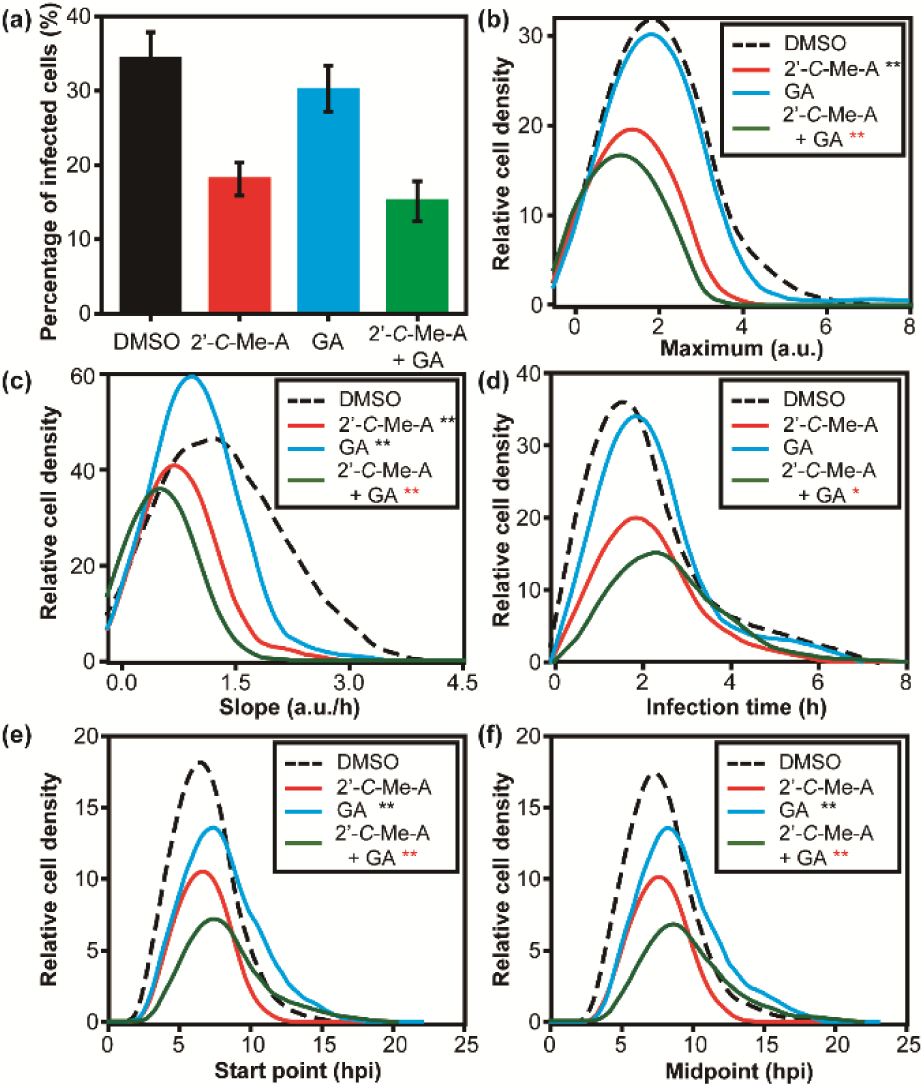
Evaluation of an antiviral drug combination: 2’-*C*-Me-A and GA. (**a**) Combination of 2’-*C*-Me-A and GA is not synergistic based on the fraction of established infections. Percentage of single, infected (green) cells was determined in the presence of 50 µM 2’-*C*-Me-A, 25 nM GA, or the combination of the two drugs. (**b**-**f**) Distributions for each parameter in the presence of the combination were compared to that in the presence of GA alone using a t-test. Numerical values for experimental parameters are provided in **Table S9** and statistical analysis in **Table S10**. The parameters presented in the panels are as follows: (**b**) maximum; (**c**) slope; (**d**) infection time; (**e**) start point; (**f**) midpoint.

### Evaluation of single-cell data by using principal component analysis (PCA)

Our evaluation of three classes of anti-PV drugs revealed three unique signatures based on changes to the phenomenological parameters used to describe infection dynamics. We reasoned that PCA might provide an even more robust approach to compare datasets using our five parameters. As shown in **Fig. 5**, PCA resolves each class of inhibitor from the other, as well as from outcomes in the absence of drug. Importantly, the mechanistically related but chemically distinct inhibitors of HSP90 cluster by PCA (see GA and GS in **Fig. 5**). We conclude that single-cell analysis of antiviral therapeutics can provide not only information on the efficacy of a drug but also information on the mechanism of action. This experimental paradigm should have a transformative impact on the drug-development process.

**Figure 5.**
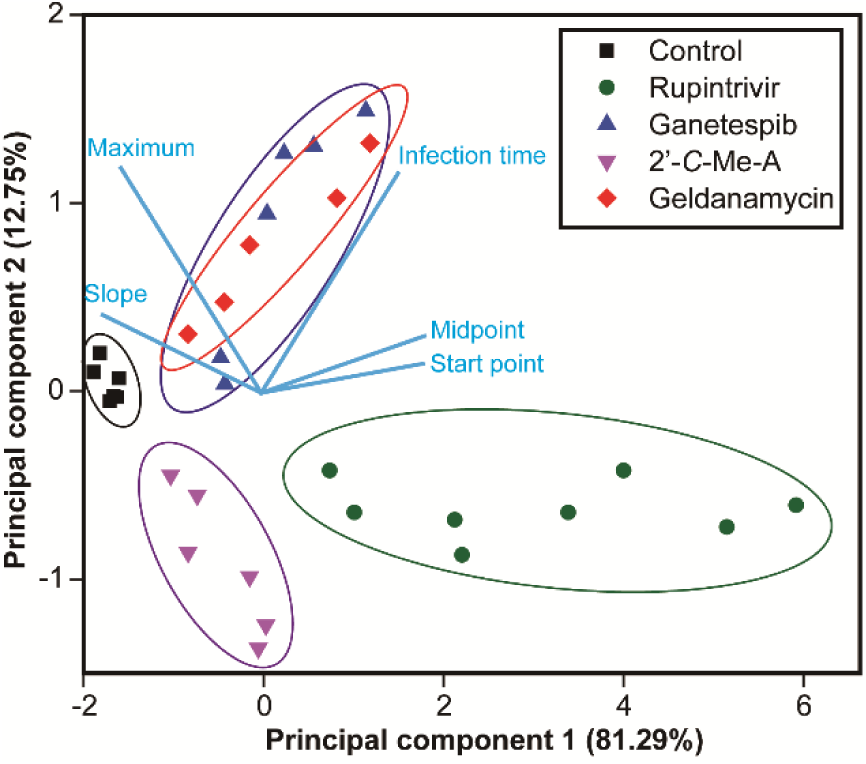
Mechanistic classes of antiviral agents distinguished by principal component analysis (PCA) of single-cell data. Points representing groups of single cells are colored according to compound identities with each point representing a different drug-treatment concentration. PCA was performed using the mean values of the maximum, slope, infection time, start point and midpoint from the various drug-treatment concentrations and control groups. The top two principal components accounted for 94% of the total variance. The light blue lines indicate the relationship between variables in the space of the first two components.

## DISCUSSION

The era of single-cell virology has begun (Combe et al., 2015; Guo et al., 2017; Russell et al., 2018). Infection outcomes in cells are determined, in part, by stochastic processes (Heldt et al., 2015). For example, the presence and abundance of viral-restriction factors and viral-replication-promoting factors of the host are largely governed by stochastic gene expression (Raj and van Oudenaarden, 2008). As a result, substantial between-cell variation exists in all quantifiable parameters of infection. An underappreciated consequence of the behavior of systems such as these is the non-self-averaging nature of these systems, which means that the population average of any single-cell parameter will almost certainly fail to represent the dynamics within any single cell (Derrida, 1997; Wiseman and Domany, 1998). The practical complication arising from this phenomenon is that changes in the average kinetics observed in a growth curve for a virus caused by a perturbation, for example exposure to an antiviral therapeutic, will not inform accurately the step(s) of the viral lifecycle impacted most by the perturbation.

With this perspective in mind, it is not surprising that our previous evaluation of an antiviral ribonucleotide using a single-cell approach revealed the unexpected finding that the most-fit members of the viral population were most susceptible to this class of compounds (Guo et al., 2017). This observation left one major unresolved question: do all effective antiviral therapeutics target the same subpopulation of viruses or do unique signatures exist for distinct mechanistic classes of antiviral therapeutics? Addressing these questions is the central objective of this study.

Our first-generation microfluidic device for on-chip investigation of viral infection dynamics relied on cell density as the mechanism to achieve isolation of single cells in wells of the device, an approach which leaves most wells of the device empty (Guo et al., 2017). We have redesigned the device to add physical cell-trapping structures to each well. The outcome of which is the ability to achieve single-cell occupancy of ~90% of the ~6000 wells of the device (**Figs. 1, S1**-**S3**). The enhanced occupancy enabled the complete characterization of antiviral therapeutics by permitting the acquisition of complete dose-response curves (e.g. **Fig. 2a**).

The three classes of antiviral therapeutics chosen target a viral polymerase (2’-C-methyladenosine, 2’-*C*-Me-A, **Fig. S4**), a viral protease (rupintrivir, **Fig. 2**), or a host factor, HSP90, (geldanamycin, GA, **Fig. 3**; ganetespib, GS, **Fig. S6**). Our data-analysis pipeline emphasizes five phenomenological parameters with correlates to traditional parameters for assessment of the viral lifecycle.^7^ The major finding of this study is that each class of antiviral therapeutics exhibited a unique signature with respect to the five phenomenological parameters measured and so much so that it was possible to use principal component analysis (PCA) to stratify the different therapeutic classes (**Fig. 5**). Importantly, both GA and GS overlapped by PCA despite their substantial differences in structure and efficacy (**Fig. 5**).

To our knowledge, this study represents the first analysis of a range of antiviral therapeutics on viral infection dynamics at the single-cell level. The resolution afforded by this approach is unprecedented when compared to other cell-based approaches, which only provide a measure of efficacy. Given the substantial effort required to go from compound to mechanism using the traditional experimental paradigm, the ability of single-cell analysis to inform mechanism should make this approach a welcome addition to the drug discovery and development toolbox. It is not uncommon for analogues to be synthesized during the drug-development process that lose specificity or even function by a different mechanism of action, single-cell analysis has the potential to reveal changes such as these at the start of the analysis instead of much, much later in the development process. Finally, although we have used antiviral agents to demonstrate the power of single-cell analysis, use of this technology and approach will be applicable to the discovery and development of any class of therapeutics that can be assessed in cell culture.

## METHODS

### Cells

HeLa and HeLa S3 cells were obtained from American Type Culture Collection (ATCC) and maintained in DMEM/F12 (1:1) (Life Technologies) supplemented with 10% fetal bovine serum (FBS, Atlanta Biologicals) and 100 IU/mL penicillin–streptomycin (Corning), in a humidified atmosphere of 95% air and 5% CO_2_ at 37 °C.

### Viruses

To generate EGFP-tagged poliovirus used in this work, pMo-EGFP-PV-WT plasmid was linearized with ApaI and purified with QIAEX II Gel Extraction Kit (Qiagen, Netherlands). With the linearized plasmid as the template, viral RNA was transcribed at 37°C for 5.5 hours in a 20-μL reaction medium containing 350 mM HEPES pH 7.5, 32 mM magnesium acetate, 40 mM dithiothreitol (DTT), 2 mM spermidine, 28 mM nucleoside triphosphates (NTPs), 0.025 µg/µL linearized DNA, and 0.025 µg/µL T7 RNA polymerase. After removing magnesium pyrophosphate in the mixture by centrifugation for 2 minutes, RNA concentration was measured by scanning the gel at fluorescence mode with a Typhoon 8600 scanner (Promega, USA). Then, 5 µg of viral RNA were utilized to transfect HeLa cells cultured at 37°C by electroporation. Virus was harvested by three repeated freeze-thaw cycles, centrifuged at 3000 rpm for 5 minutes, and suspended in 0.5% nonidet P-40 (NP-40). For purification, the supernatant was mixed with 1 volume of 20% PEG-8000/1 M NaCl solution and incubated overnight at 4°C. After centrifugation at 8000 × g for 10 minutes at 4°C, the pellet was resuspended in PBS and filtered with Centricon® Plus-70 (EMD Millipore, USA). Plaque assay was performed to determine the virus titer.

### Microfluidic devices

The microfluidic layers were fabricated from polydimethylsiloxane (PDMS, GE RTV615) by standard soft lithography techniques. The molds were fabricated by coating photoresist SU-8 25/50 with desired thicknesses on a silicon wafer and photolithographic patterning (Figure S1 and S2). Premixed PDMS prepolymer and curing agent (ratio 5/1) were poured onto the mold for valve layer and cured. Premixed PDMS prepolymer and curing agent (ratio 20/1) were spin-coated on the mold for channel layer at 1500 rpm for 1 minute and incubated at 65 °C for 20 min. Then the valve layer was released from the mold and placed on the channel layer with alignment. The assembly was baked at 65 °C overnight for efficient bonding. The assembly was further bonded with alignment to the microwells layer cured from premixed PDMS prepolymer and curing agent (ratio 20/1) by baking at 65 °C overnight. The obtained device was further bonded to a glass slide to prevent deformation when applying a pressure to the valves.

### On-chip experiments

Dimethyl sulfoxide (DMSO) and rupintrivir were purchased from Sigma Chemical Co. (St. Louis, MO, USA). 2’-*C*-methyladenosine (2’-*C*-Me-A) was obtained from Carbosynth Limited (Compton, Berkshire, UK). Solutions of geldanamycin and ganetespib in DMSO were kindly provided by Dr. Xin Zhang (Department of Chemistry, The Pennsylvania State University) and Dr. Judith Frydman (Department of Biology, Stanford University), respectively. All the antiviral compounds were pre-diluted in DMSO for use. For no drug groups, same volumes of DMSO were spiked into the culture medium.

For each group, 2 × 10^5^ HeLa S3 cells were cultured with drug-containing medium for 1 h. Then the cells were centrifuged, resuspended in PBS, and mixed with EGFP-PV at the MOI of 0.5 PFU/cell. After shaking at 140 rpm for 30 min, the cells were centrifugated and resuspended in mediums with corresponding treatments. With a PHD Ultra syringe pump (Harvard Apparatus), the cell suspensions were infused to the inlets of a microfluidic device for trapping of single-cells for 5 min. Afterwards, a pressure of 30 psi was applied to the valves to isolate the microwells.

The device was placed in the chamber of a stage top WSKM GM2000 incubation system (Tokai, Japan) adapted to a Nikon Eclipse Ti inverted microscope (Nikon, Japan) equipped with a ProScan II motorized flat top stage (Prior Scientific, USA). With this setup, bright-field and fluorescence images of the microwells were automatically acquired every 30 minutes from 2.5 hpi to 24 hpi. In each image, 36 (6 × 6) microwells were included with a CFI60 Plan Apochromat Lambda 10× objective and a Hamamatsu C11440 camera.

### Data processing

A customized MATLAB script was employed to extract the fluorescence intensity and background intensity for each microwell in each fluorescent image. (Fluorescence intensity - Background)/Background was calculated to represent the relative intensity. Meanwhile, wells containing two or more infected cells and showing auto-fluorescence or out-of-focus signals were manually excluded. As a result, the fluorescence intensity of each infected single-cell over time could be obtained. With a customized package of R, the maximum, slope, infection time, start point and midpoint were derived (Caglar et al., 2018). For comparison of different experiments, the intensities were further normalized to uniformize the mean values of the maximums for the no drug groups (2.00 a.u. in this work). The distributions of cell densities were plotted with the area under each curve representing the relative number of infected cells (given 100 cells for no drug group).

## Supplementary Information

The following is provided: design of the microfluidic device; influence of channel width on single-cell trapping; effect of 2’-*C*-meA and ganetespib at different concentrations on viral replication; means, standard deviations and p-values of all of the parameters measured for all of the experiments.

## Acknowledgements

This work was supported by grant AI12056 from NIAID, NIH to C.O.W. and C.E.C.. A.W. is the recipient of a postdoctoral fellowship from the American Heart Association. W.L. thanks Dr. Sixing Li, Dr. Peng Li, Dr. Po-Hsun Huang and Liqiang Ren for helpful discussions. C.E.C. thanks the following: Dr. Susanna Manrubia for sharing her thoughts on non-self-averaging systems; Dr. Judith Frydman and Dr. Raul Andino for encouraging us to evaluate inhibitors of HSP90.

## Author Contributions

W.L. and C.E.C. designed the experiments. W.L. conducted the experiments. A.W and J.J.A. contributed unique reagents. W.L., M.U.C., and Z.M. performed image and data analysis. W.L., M.U.C., A.W., J.J.A., C.O.W., and C.E.C analyzed the data. W.L. and C.E.C wrote the manuscript.

## Graphical Abstract

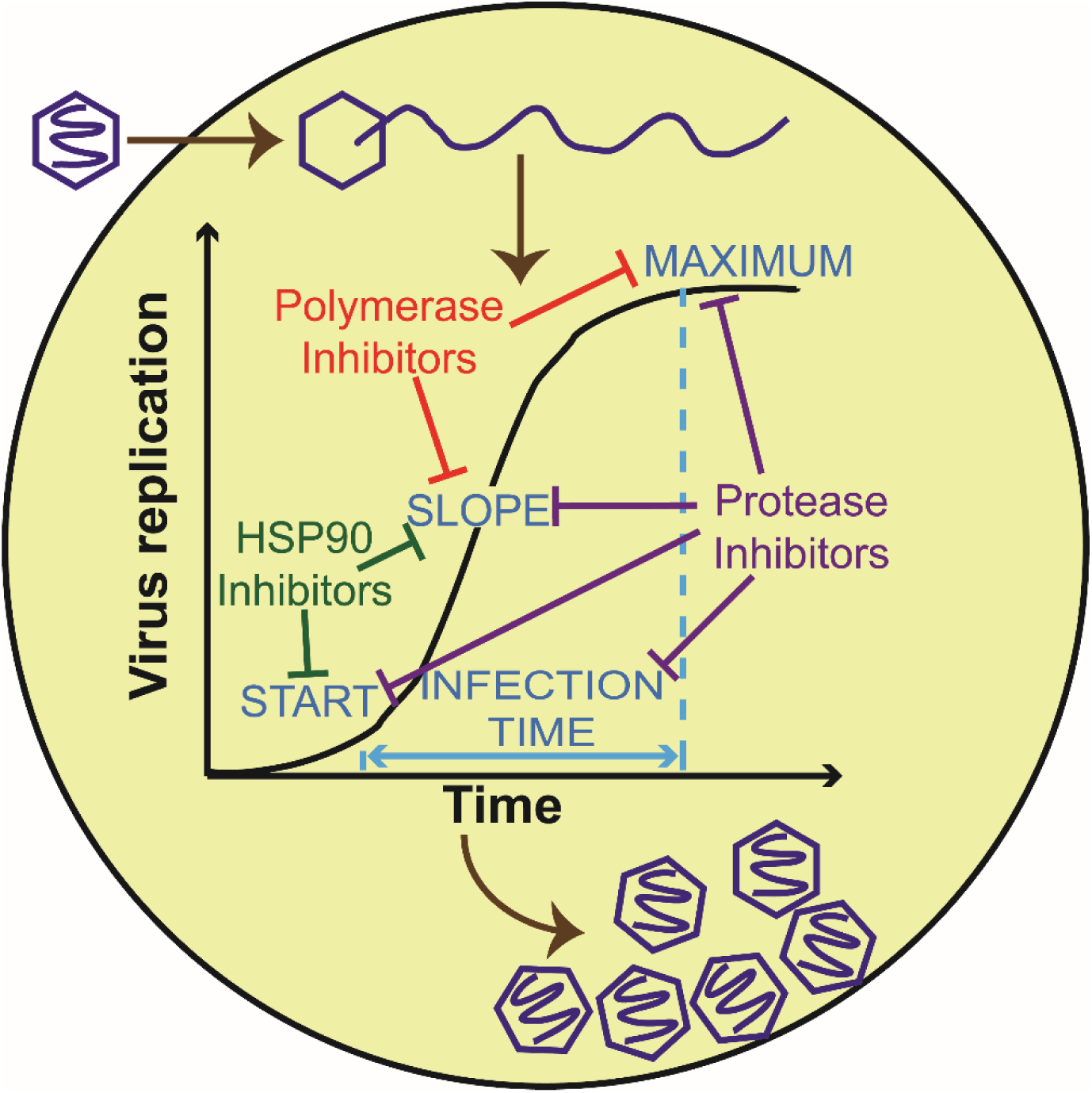

