## Supplementary Information for "More than efficacy revealed by single-cell analysis of antiviral therapeutics"

### Table of content:

**Figure S1** A microfluidic device containing cell-trapping microstructures for high-througput analysis of single-cell infections. (S2)

**Figure S2** Cross-section of the microfluidic device. (S3)

**Figure S3** Optimization of single-cell trapping in microfluidic wells. (S4)

**Figure S4** Evaluation of 2’-*C*-methyl-adenosine nucleoside (2’-*C*-Me-A), a PV RdRp inhibitor. (S5)

**Figure S5** Translation of eGFP mRNA and activation of fluorophore are normal in the presence of GA and GS. (S6)

**Figure S6** Evaluation of ganetespib (GS), an HSP90 inhibitor. (S7)

**Figure S7** Evaluation of an antiviral drug combination: 2’-*C*-Me-A and GS. (S8)

**Table S1** Means and standard deviations for the 2’-*C*-Me-A experiment. (S9)

**Table S2** P-values between groups based on t-test for the 2’-*C*-Me-A experiment. (S9)

**Table S3** Means and standard deviations for the rupintrivir experiment. (S10)

**Table S4** P-values between groups based on t-test for the rupintrivir experiment. (S10)

**Table S5** Means and standard deviations for the geldanamycin experiment. (S11)

**Table S6** P-values between groups based on t-test for the geldanamycin experiment. (S11)

**Table S7** Means and standard deviations for the ganetespib experiment. (S12)

**Table S8** P-values between groups based on t-test for the ganetespib experiment. (S12)

**Table S9** Means and standard deviations for antiviral synergy of 2’-*C*-Me-A and GA. (S13)

**Table S10** P-values between groups based on t-test for antiviral synergy of 2’-*C*-Me-A and GA. (S13)

**Table S11** Means and standard deviations for antiviral synergy of 2’-*C*-Me-A and GS. (S14)

**Table S12** P-values between groups based on t-test for antiviral synergy of 2’-*C*-Me-A and GS. (S14)

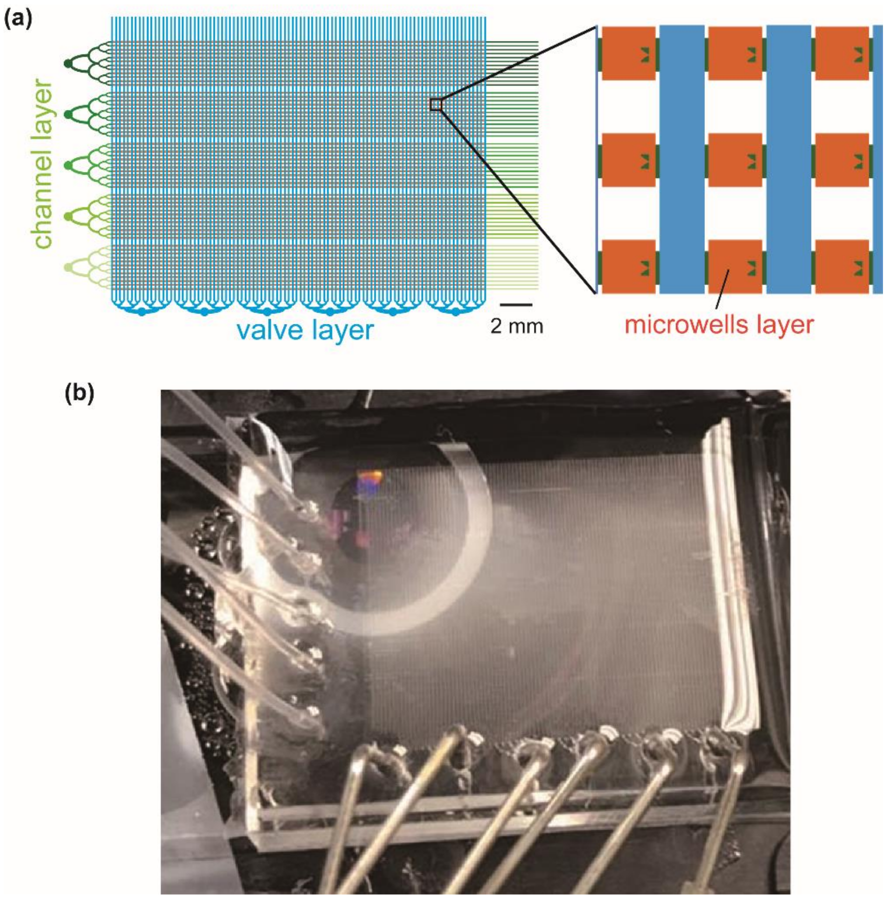


**Figure S1. A microfluidic device containing cell-trapping microstructures for high-througput analysis of single-cell infections.** (**a**) Organization of microfluidic device into five sections of 1140 (12×95) microwells permits five independent experiment to be performed on each device. (**b**) Shown below is an image of a device mounted on the stage of the microscope.

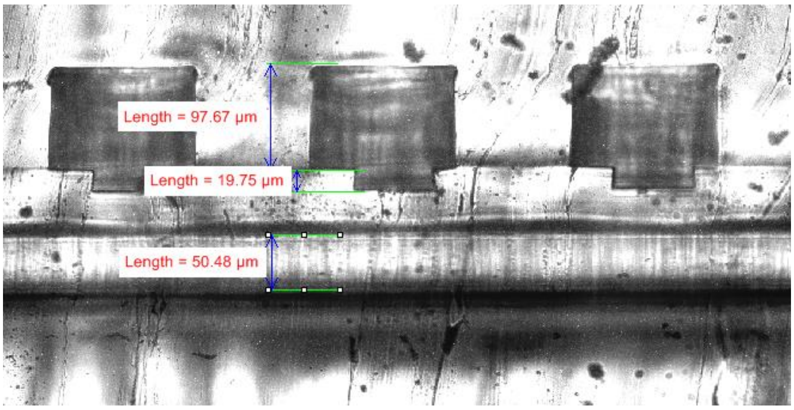


**Figure S2. Cross-section of the microfluidic device.** The microwells for cell culture, channels for cell loading, and valve channels for isolation of the microwells are ~100 µm, ~20 µm, and ~50 µm deep, respectively.

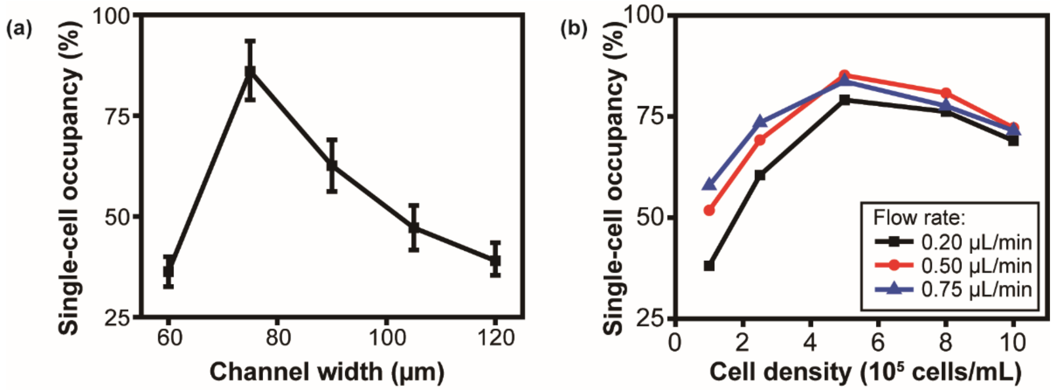


**Figure S3. Optimization of single-cell trapping in microfluidic wells.** (**a**) Influence of channel width on single-cell trapping efficiency. Cells ($5 \times 10^5$ cells/mL)were infused into the device at a flow rate of 0.5 µL/min. When the channel width was 60 µm, the gap between the trapping structure and the sidewall of the channel was so narrow that more than one cell was trapped in most wells. (**b**) Influence of cell density on the single-cell occupancy at different infusion rates. Channel width was 75 µm.

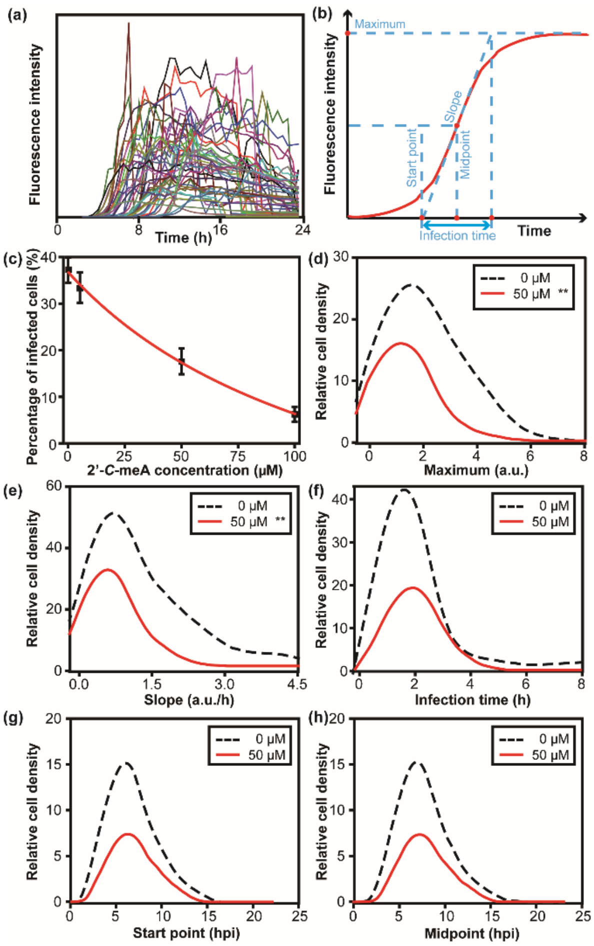


**Figure S4**. **Evaluation of 2'-*C*-methyl-adenosine nucleoside (2'-*C*-Me-A), a PV RdRp inhibitor.** (**a**) Between-cell variability of PV replication. HeLa S3 cells were infected with PV-eGFP at a MOI of 0.5 PFU/cell, loaded on the microfluidic device, and observed for green fluorescence every 30 min for a 24 h. (**b**) Dose-response analysis. Percentage of single, infected (green) cells was determined as a function of 2'-*C*-Me-A concentration. (**c**) Data analysis. Each growth curve can be described minimally using the parameters indicated. The distribution for each parameter is used to determine the impact of an experimental variable on the experimental outcome. (**d-h**) Distributions for each parameter in the presence of 50 μM 2'-*C*-Me-A were compared to that in the absence of drug using a t-test. A single asterisk in the key indicates a p-value less than 0.05; two asterisks indicate a p-value less than 0.005. Numerical values for experiemental parameters are provided in **Table S1** and statistical analysis in **Table S2**. The parameters presented in the panels are as follows: (**d**) maximum; (**e**) slope; (**f**) infection time; (**g**) start point; (**h**) midpoint.

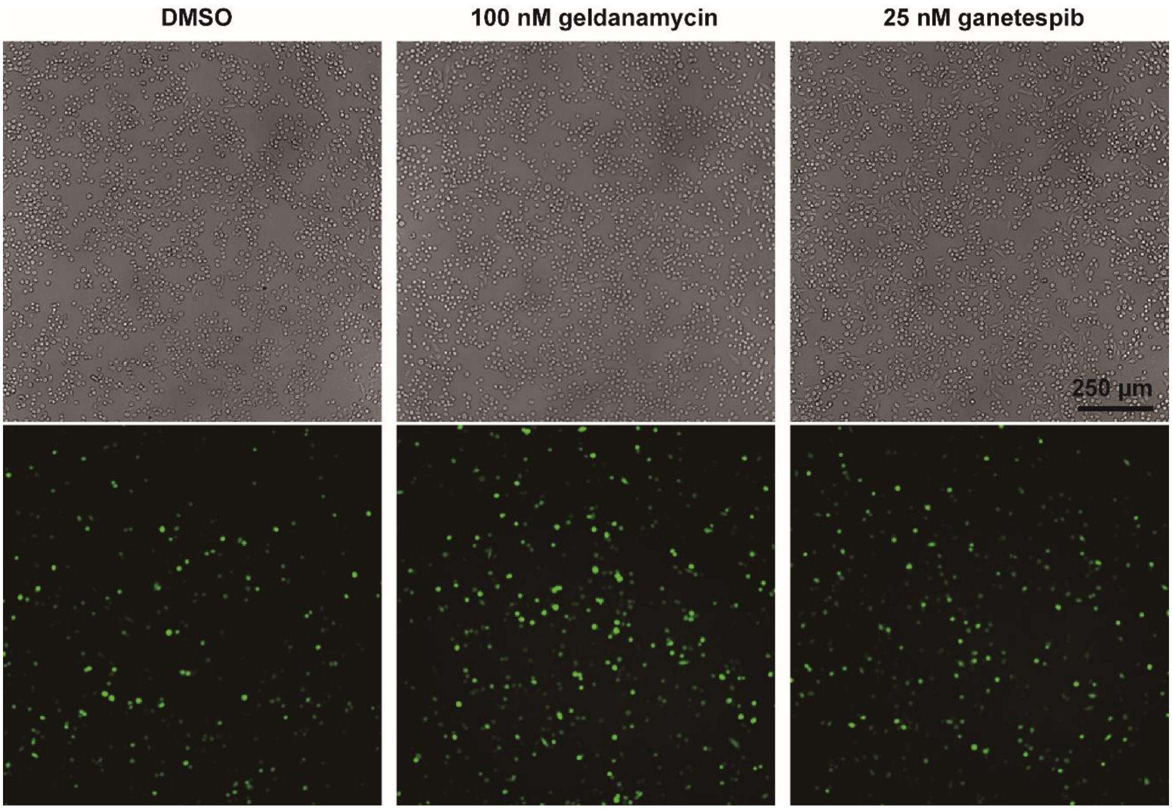


**Figure S5**. **Translation of eGFP mRNA and activation of fluorophore are normal in the presence of GA and GS.** GFP RNA was transfected into HeLa S3 cells and the expression of GFP was evaluated in the presence of GA or GS at the highest concentration used in the experiments described herein. Fluorescent images taken at 4 hours post-transfection exhibited similar levels of GFP expression as control (DMSO) group.

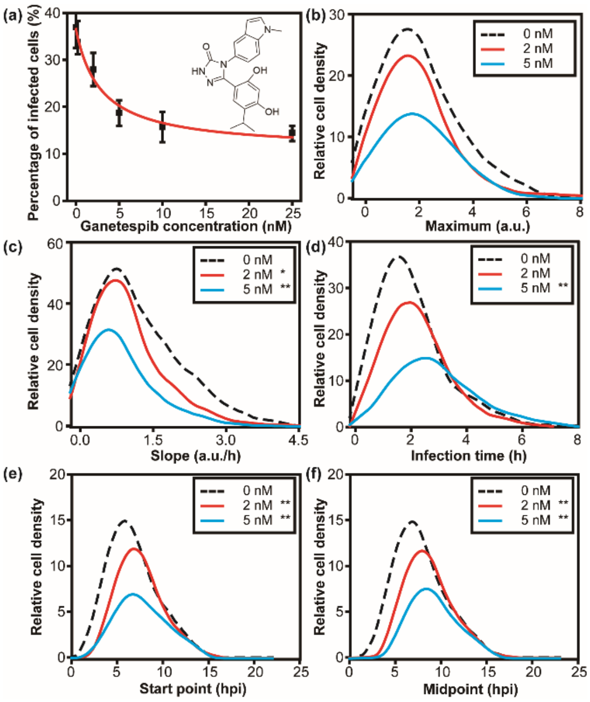


**Figure S6**. **Evaluation of ganetespib (GS), an HSP90 inhibitor.** (**a**) Dose-response analysis. Percentage of single, infected (green) cells was determined as a function of GS concentration. (**b-f**) Distributions for each parameter in the presence of 2 or 5 nM GS were compared to that in the absence of drug using a t-test. A single asterisk in the key indicates a p-value less than 0.05; two asterisks indicate a p-value less than 0.005. Numerical values for experimental parameters are provided in **Table S7** and statistical analysis in **Table S8**. The parameters presented in the panels are as follows: (**b**) maximum; (**c**) slope; (**d**) infection time; (**e**) start point; (**f**) midpoint.

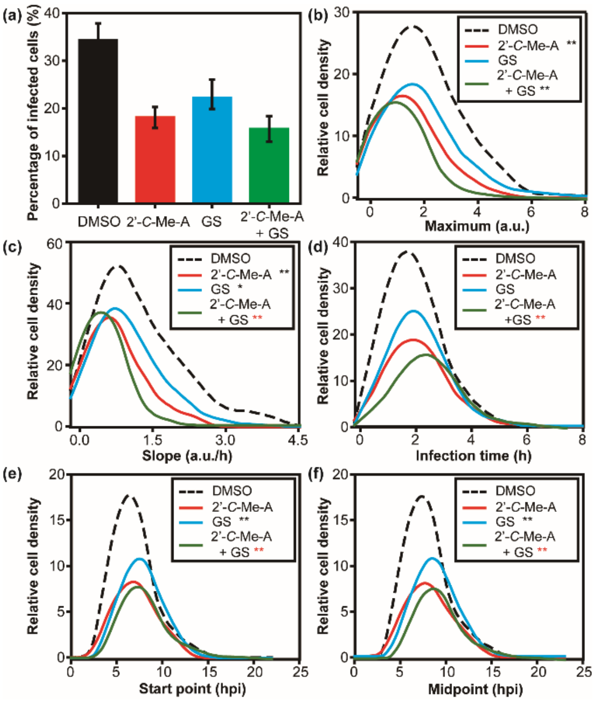


**Figure S7. Evaluation of an antiviral drug combination: 2'-*C*-Me-A and GS.** (**a**) Percentage of single, infected cells was determined in the presence of 50 μM 2'-C-Me-A, 2 nM GS, or the combination of the two drugs. The control (DMSO) had 34 ± 4% cells infected. This values was lowered to 18 ± 2%, 22 ± 3%, and 16 ± 3% in the presence of 2'-*C*-Me-A, GS, or the combination, respectively. (**b-f**) Distributions for each parameter under the various conditions were compared using a t-test. Numerical values for experimental parameters are provided in **Table S11** and statistical analysis in **Table S12**. The parameters presented in the panels are as follows: (**b**) maximum; (**c**) slope; (**d**) infection time; (**e**) start point; (**f**) midpoint.

**Table S1** Means and standard deviations for the 2'-*C*-Me-A experiment.

| Group | Cell Number | Maximum (a.u.) | | Slope (a.u./h) | | Infection time (h) | | Start point (hpi) | | Midpoint (hpi) | |
|---|---|---|---|---|---|---|---|---|---|---|---|
| | | Mean | SD | Mean | SD | Mean | SD | Mean | SD | Mean | SD |
| **0 µM** | 206 | 2.00 | 1.34 | 1.17 | 0.91 | 2.00 | 1.05 | 6.65 | 2.34 | 7.65 | 2.37 |
| **5 µM** | 76 | 1.75 | 1.24 | 1.01 | 0.88 | 2.08 | 0.79 | 6.77 | 2.03 | 7.81 | 2.13 |
| **50 µM** | 91 | 1.32 | 0.84 | 0.70 | 0.46 | 2.11 | 0.72 | 6.78 | 2.02 | 7.84 | 2.06 |

**Table S2** P-values between groups based on t-test for the 2'-*C*-Me-A experiment.

| | Group | 0 µM | 5 µM | 50 µM |
|---|---|---|---|---|
| **Maximum** | 0 µM | 1 | 0.162897 | **0.000011** |
| | 5 µM | 0.162897 | 1 | **0.008076** |
| | 50 µM | **0.000011** | **0.008076** | 1 |
| **Slope** | Group | 0 µM | 5 µM | 50 µM |
| | 0 µM | 1 | 0.188607 | **< 0.00001** |
| | 5 µM | 0.188607 | 1 | **0.003502** |
| | 50 µM | **< 0.00001** | **0.003502** | 1 |
| **Infection time** | Group | 0 µM | 5 µM | 50 µM |
| | 0 µM | 1 | 0.550896 | 0.341086 |
| | 5 µM | 0.550896 | 1 | 0.75737 |
| | 50 µM | 0.341086 | 0.75737 | 1 |
| **Start point** | Group | 0 µM | 5 µM | 50 µM |
| | 0 µM | 1 | 0.689734 | 0.649419 |
| | 5 µM | 0.689734 | 1 | 0.981178 |
| | 50 µM | 0.649419 | 0.981178 | 1 |
| **Midpoint** | Group | 0 µM | 5 µM | 50 µM |
| | 0 µM | 1 | 0.604706 | 0.517508 |
| | 5 µM | 0.604706 | 1 | 0.937393 |
| | 50 µM | 0.517508 | 0.937393 | 1 |

**Table S3** Means and standard deviations for the rupintrivir experiment.

| Group | Cell Number | Maximum (a.u.) | | Slope (a.u./h) | | Infection time (h) | | Start point (hpi) | | Midpoint (hpi) | |
|---|---|---|---|---|---|---|---|---|---|---|---|
| | | Mean | SD | Mean | SD | Mean | SD | Mean | SD | Mean | SD |
| **0 nM** | 213 | 2.00 | 1.42 | 1.18 | 0.93 | 2.06 | 1.15 | 6.73 | 2.08 | 7.76 | 2.15 |
| **10 nM** | 124 | 1.35 | 0.83 | 0.61 | 0.40 | 2.48 | 1.33 | 7.60 | 2.48 | 8.84 | 2.53 |
| **20 nM** | 117 | 1.09 | 0.79 | 0.49 | 0.35 | 2.56 | 1.48 | 8.58 | 2.45 | 9.86 | 2.66 |
| **50 nM** | 32 | 0.57 | 0.38 | 0.20 | 0.13 | 3.21 | 1.38 | 12.36 | 2.58 | 13.97 | 2.39 |

**Table S4** P-values between groups based on t-test for the rupintrivir experiment.

| | Group | 0 nM | 10 nM | 20 nM | 50 nM |
|---|---|---|---|---|---|
| **Maximum** | 0 nM | 1 | **0.000019** | **< 0.00001** | **< 0.00001** |
| | 10 nM | **0.000019** | 1 | **0.013749** | **< 0.00001** |
| | 20 nM | **< 0.00001** | **0.013749** | 1 | **0.000477** |
| | 50 nM | **< 0.00001** | **< 0.00001** | **0.000477** | 1 |
| **Slope** | Group | 0 nM | 10 nM | 20 nM | 50 nM |
| | 0 nM | 1 | **< 0.00001** | **< 0.00001** | **< 0.00001** |
| | 10 nM | **< 0.00001** | 1 | **0.011982** | **< 0.00001** |
| | 20 nM | **< 0.00001** | **0.011982** | 1 | **< 0.00001** |
| | 50 nM | **< 0.00001** | **< 0.00001** | **< 0.00001** | 1 |
| **Infection time** | Group | 0 nM | 10 nM | 20 nM | 50 nM |
| | 0 nM | 1 | **0.010999** | **0.005387** | **< 0.00001** |
| | 10 nM | **0.010999** | 1 | 0.673427 | **0.006574** |
| | 20 nM | **0.005387** | 0.673427 | 1 | **0.025759** |
| | 50 nM | **< 0.00001** | **0.006574** | **0.025759** | 1 |
| **Start point** | Group | 0 nM | 10 nM | 20 nM | 50 nM |
| | 0 nM | 1 | **0.003677** | **0** | **< 0.00001** |
| | 10 nM | **0.003677** | 1 | **0.002288** | **< 0.00001** |
| | 20 nM | **< 0.00001** | **0.002288** | 1 | **< 0.00001** |
| | 50 nM | **< 0.00001** | **< 0.00001** | **< 0.00001** | 1 |
| **Midpoint** | Group | 0 nM | 10 nM | 20 nM | 50 nM |
| | 0 nM | 1 | **0.0005** | **0** | **< 0.00001** |
| | 10 nM | **0.0005** | 1 | **0.002627** | **< 0.00001** |
| | 20 nM | **< 0.00001** | **0.002627** | 1 | **< 0.00001** |
| | 50 nM | **< 0.00001** | **< 0.00001** | **< 0.00001** | 1 |

**Table S5** Means and standard deviations for the geldanamycin (GA) experiment.

| Group | Cell Number | Maximum (a.u.) | | Slope (a.u./h) | | Infection time (h) | | Start point (hpi) | | Midpoint (hpi) | |
|---|---|---|---|---|---|---|---|---|---|---|---|
| | | Mean | SD | Mean | SD | Mean | SD | Mean | SD | Mean | SD |
| **0 nM** | 130 | 2.00 | 0.99 | 1.28 | 0.78 | 2.11 | 1.30 | 6.57 | 2.45 | 7.63 | 2.78 |
| **10 nM** | 90 | 1.91 | 0.89 | 1.17 | 0.73 | 2.06 | 1.14 | 6.44 | 2.33 | 7.47 | 2.62 |
| **25 nM** | 130 | 2.00 | 0.95 | 1.00 | 0.50 | 2.30 | 1.08 | 7.65 | 2.96 | 8.80 | 3.17 |
| **100 nM** | 86 | 1.84 | 0.77 | 0.77 | 0.54 | 3.06 | 1.37 | 8.03 | 3.11 | 9.56 | 3.33 |

**Table S6** P-values between groups based on t-test for the geldanamycin (GA) experiment.

| | | | | | |
|---|---|---|---|---|---|
| **Maximum** | Group | 0 nM | 10 nM | 25 nM | 100 nM |
| | 0 nM | 1 | 0.510088 | 0.98185 | 0.191109 |
| | 10 nM | 0.510088 | 1 | 0.55277 | 0.569293 |
| | 25 nM | 0.98185 | 0.55277 | 1 | 0.225501 |
| | 100 nM | 0.191109 | 0.569293 | 0.225501 | 1 |
| **Slope** | Group | 0 nM | 10 nM | 25 nM | 100 nM |
| | 0 nM | 1 | 0.317362 | **0.004012** | **< 0.00001** |
| | 10 nM | 0.317362 | 1 | 0.085059 | **0.000103** |
| | 25 nM | **0.004012** | 0.085059 | 1 | **0.004901** |
| | 100 nM | **< 0.00001** | **0.000103** | **0.004901** | 1 |
| **Infection time** | Group | 0 nM | 10 nM | 25 nM | 100 nM |
| | 0 nM | 1 | 0.6764 | 0.294167 | **< 0.00001** |
| | 10 nM | 0.6764 | 1 | 0.150681 | **< 0.00001** |
| | 25 nM | 0.294167 | 0.150681 | 1 | **0.000276** |
| | 100 nM | **< 0.00001** | **< 0.00001** | **0.000276** | 1 |
| **Start point** | Group | 0 nM | 10 nM | 25 nM | 100 nM |
| | 0 nM | 1 | 0.336891 | **0.004456** | **0.000162** |
| | 10 nM | 0.336891 | 1 | **0.001152** | **0.000064** |
| | 25 nM | **0.004456** | **0.001152** | 1 | 0.420694 |
| | 100 nM | **0.000162** | **0.000064** | 0.420694 | 1 |
| **Midpoint** | Group | 0 nM | 10 nM | 25 nM | 100 nM |
| | 0 nM | 1 | 0.350551 | **0.005621** | **0.000012** |
| | 10 nM | 0.350551 | 1 | **0.001251** | **< 0.00001** |
| | 25 nM | **0.005621** | **0.001252** | 1 | 0.152652 |
| | 100 nM | **0.000012** | **< 0.00001** | 0.152652 | 1 |

**Table S7** Means and standard deviations for the ganetespib (GS) experiment.

| Group | Cell Number | Maximum (a.u.) | | Slope (a.u./h) | | Infection time (h) | | Start point (hpi) | | Midpoint (hpi) | |
|---|---|---|---|---|---|---|---|---|---|---|---|
| | | Mean | SD | Mean | SD | Mean | SD | Mean | SD | Mean | SD |
| **0 nM** | 222 | 2.00 | 1.45 | 1.17 | 0.82 | 2.01 | 1.09 | 6.70 | 2.37 | 7.70 | 2.44 |
| **2 nM** | 155 | 1.92 | 1.33 | 0.97 | 0.66 | 2.23 | 1.02 | 7.51 | 2.16 | 8.63 | 2.21 |
| **5 nM** | 198 | 1.97 | 1.17 | 0.84 | 0.67 | 2.92 | 1.35 | 7.69 | 2.46 | 9.15 | 2.20 |
| **25 nM** | 136 | 1.92 | 0.94 | 0.74 | 0.39 | 3.11 | 1.68 | 7.93 | 2.32 | 9.48 | 2.55 |

**Table S8** P-values between groups based on t-test for the ganetespib (GS) experiment.

| | Group | 0 nM | 2 nM | 5 nM | 25 nM |
|---|---|---|---|---|---|
| **Maximum** | 0 nM | 1 | 0.609003 | 0.842967 | 0.733147 |
| | 2 nM | 0.609003 | 1 | 0.672029 | 0.993707 |
| | 5 nM | 0.842967 | 0.672029 | 1 | 0.777625 |
| | 25 nM | 0.733147 | 0.993707 | 0.777625 | 1 |
| **Slope** | Group | 0 nM | 2 nM | 5 nM | 25 nM |
| | 0 nM | 1 | **0.028514** | **0.00011** | **0.002769** |
| | 2 nM | **0.028514** | 1 | 0.061673 | **0.041778** |
| | 5 nM | **0.00011** | 0.061673 | 1 | 0.382645 |
| | 25 nM | **0.002769** | **0.041778** | 0.382645 | 1 |
| **Infection time** | Group | 0 nM | 2 nM | 5 nM | 25 nM |
| | 0 nM | 1 | 0.08553 | **< 0.00001** | **< 0.00001** |
| | 2 nM | 0.08553 | 1 | **< 0.00001** | **< 0.00001** |
| | 5 nM | **< 0.00001** | **<0.00001** | 1 | 0.461503 |
| | 25 nM | **<0.00001** | **< 0.00001** | 0.461503 | 1 |
| **Start point** | Group | 0 nM | 2 nM | 5 nM | 25 nM |
| | 0 nM | 1 | **0.003242** | **0.000419** | **0.000743** |
| | 2 nM | **0.003242** | 1 | 0.462347 | 0.306492 |
| | 5 nM | **0.000419** | 0.462347 | 1 | 0.600001 |
| | 25 nM | **0.000743** | 0.306492 | 0.600001 | 1 |
| **Midpoint** | Group | 0 nM | 2 nM | 5 nM | 25 nM |
| | 0 nM | 1 | **0.00114** | **< 0.00001** | **< 0.00001** |
| | 2 nM | **0.00114** | 1 | **0.026382** | **0.044117** |
| | 5 nM | **< 0.00001** | **0.026382** | 1 | 0.42665 |
| | 25 nM | **< 0.00001** | **0.044117** | 0.42665 | 1 |

**Table S9** Means and standard deviations for antiviral synergy of 2’-*C*-Me-A and GA.

| Group | Cell Number | Maximum (a.u.) | | Slope (a.u./h) | | Infection time (h) | | Start point (hpi) | | Midpoint (hpi) | |
|---|---|---|---|---|---|---|---|---|---|---|---|
| | | Mean | SD | Mean | SD | Mean | SD | Mean | SD | Mean | SD |
| **DMSO** | 145 | 2.00 | 0.94 | 1.29 | 0.75 | 2.04 | 1.26 | 6.71 | 1.65 | 7.73 | 1.96 |
| **2’-*C*-Me-A** | 86 | 1.37 | 0.48 | 0.76 | 0.39 | 2.09 | 0.93 | 6.62 | 1.08 | 7.67 | 1.26 |
| **GA** | 129 | 1.86 | 0.91 | 0.94 | 0.45 | 2.27 | 1.07 | 7.91 | 2.29 | 9.04 | 2.47 |
| **2’-*C*-Me-A + GA** | 79 | 1.10 | 0.41 | 0.51 | 0.29 | 2.55 | 1.06 | 8.21 | 2.38 | 9.48 | 2.54 |

**Table S10** P-values between groups based on t-test for antiviral synergy of 2’-*C*-Me-A and GA.

| | Group | DMSO | 2’-*C*-Me-A | GA | 2’-*C*-Me-A + GA |
|---|---|---|---|---|---|
| **Maximum** | DMSO | 1 | **< 0.00001** | 0.364767 | **< 0.00001** |
| | 2’-*C*-Me-A | **< 0.00001** | 1 | **0.000314** | **0.001179** |
| | GA | 0.364767 | **0.000314** | 1 | **< 0.00001** |
| | 2’-*C*-Me-A + GA | **< 0.00001** | **0.001179** | **< 0.00001** | 1 |
| **Slope** | Group | DMSO | 2’-*C*-Me-A | GA | 2’-*C*-Me-A + GA |
| | DMSO | 1 | **< 0.00001** | **0.000854** | **< 0.00001** |
| | 2’-*C*-Me-A | **< 0.00001** | 1 | **0.022876** | **0.000122** |
| | GA | **0.000854** | **0.022876** | 1 | **< 0.00001** |
| | 2’-*C*-Me-A + GA | **< 0.00001** | **0.000122** | **< 0.00001** | 1 |
| **Infection time** | Group | DMSO | 2’-*C*-Me-A | GA | 2’-*C*-Me-A + GA |
| | DMSO | 1 | 0.782847 | 0.225594 | **0.009664** |
| | 2’-*C*-Me-A | 0.782847 | 1 | 0.317692 | **0.013196** |
| | GA | 0.225594 | 0.317692 | 1 | 0.155885 |
| | 2’-*C*-Me-A + GA | **0.009664** | **0.013196** | 0.155885 | 1 |
| **Start point** | Group | DMSO | 2’-*C*-Me-A | GA | 2’-*C*-Me-A + GA |
| | DMSO | 1 | 0.724046 | **0.000147** | **< 0.00001** |
| | 2’-*C*-Me-A | 0.724046 | 1 | **0.000133** | **< 0.00001** |
| | GA | **0.000147** | **0.000133** | 1 | 0.479335 |
| | 2’-*C*-Me-A + GA | **< 0.00001** | **< 0.00001** | 0.479335 | 1 |
| **Midpoint** | Group | DMSO | 2’-*C*-Me-A | GA | 2’-*C*-Me-A + GA |
| | DMSO | 1 | 0.836199 | **0.000228** | **< 0.00001** |
| | 2’-*C*-Me-A | 0.836199 | 1 | **0.000184** | **< 0.00001** |
| | GA | **0.000228** | **0.000184** | 1 | 0.336234 |
| | 2’-*C*-Me-A + GA | **< 0.00001** | **< 0.00001** | 0.336234 | 1 |

**Table S11** Means and standard deviations for antiviral synergy of 2’-*C*-Me-A and GS.

| **Group** | **Cell Number** | **Maximum (a.u.)** | | **Slope (a.u./h)** | | **Infection time (h)** | | **Start point (hpi)** | | **Midpoint (hpi)** | |
|---|---|---|---|---|---|---|---|---|---|---|---|
| | | Mean | SD | Mean | SD | Mean | SD | Mean | SD | Mean | SD |
| **DMSO** | 195 | 2.00 | 1.30 | 1.20 | 0.83 | 1.97 | 0.87 | 6.90 | 1.89 | 7.88 | 1.92 |
| **2’-*C*-Me-A** | 136 | 1.36 | 0.97 | 0.74 | 0.53 | 2.09 | 0.93 | 7.10 | 1.93 | 8.14 | 2.05 |
| **GS** | 119 | 1.91 | 1.36 | 0.98 | 0.64 | 2.12 | 0.84 | 7.77 | 1.80 | 8.84 | 1.84 |
| **2’-*C*-Me-A + GS** | 87 | 1.12 | 0.97 | 0.47 | 0.31 | 2.52 | 1.06 | 7.99 | 2.13 | 9.25 | 2.19 |

**Table S12** P-values between groups based on t-test for antiviral synergy of 2’-*C*-Me-A and GS.

| | Group | DMSO | 2’-*C*-Me-A | GS | 2’-*C*-Me-A + GS |
|---|---|---|---|---|---|
| **Maximum** | DMSO | 1 | **0.000029** | 0.637994 | **< 0.00001** |
| | 2’-*C*-Me-A | **0.000029** | 1 | **0.000216** | 0.071223 |
| | GS | 0.637994 | **0.000216** | 1 | **< 0.00001** |
| | 2’-*C*-Me-A + GS | **< 0.00001** | 0.071223 | **< 0.00001** | 1 |
| **Slope** | Group | DMSO | 2’-*C*-Me-A | GS | 2’-*C*-Me-A + GS |
| | DMSO | 1 | **< 0.00001** | **0.032673** | **< 0.00001** |
| | 2’-*C*-Me-A | **< 0.00001** | 1 | **0.001144** | **0.000027** |
| | GS | **0.032673** | **0.001144** | 1 | **< 0.00001** |
| | 2’-*C*-Me-A + GS | **< 0.00001** | **0.000027** | **< 0.00001** | 1 |
| **Infection time** | Group | DMSO | 2’-*C*-Me-A | GS | 2’-*C*-Me-A + GS |
| | DMSO | 1 | 0.296104 | 0.192938 | **0.000146** |
| | 2’-*C*-Me-A | 0.296104 | 1 | 0.80909 | **0.001683** |
| | GS | 0.192938 | 0.80909 | 1 | **0.002812** |
| | 2’-*C*-Me-A + GS | **0.000146** | **0.001683** | **0.002812** | 1 |
| **Start point** | Group | DMSO | 2’-*C*-Me-A | GS | 2’-*C*-Me-A + GS |
| | DMSO | 1 | 0.431104 | **0.00062** | **0.000322** |
| | 2’-*C*-Me-A | 0.431104 | 1 | **0.004163** | **0.001395** |
| | GS | **0.00062** | **0.004163** | 1 | 0.433485 |
| | 2’-*C*-Me-A + GS | **0.000322** | **0.001395** | 0.433485 | 1 |
| **Midpoint** | Group | DMSO | 2’-*C*-Me-A | GS | 2’-*C*-Me-A + GS |
| | DMSO | 1 | 0.322303 | **0.000268** | **0.000012** |
| | 2’-*C*-Me-A | 0.322303 | 1 | **0.005235** | **0.000166** |
| | GS | **0.000268** | **0.005235** | 1 | 0.139701 |
| | 2’-*C*-Me-A + GS | **0.000012** | **0.000166** | 0.139701 | 1 |
